# Computational Repurposing and Phytochemical Screening of Inhibitors Against ClpK from *Klebsiella pneumoniae*

**DOI:** 10.1101/2025.11.28.691150

**Authors:** Tehrim Ballim, Thandeka Khoza

**Affiliations:** Department of Biochemistry, School of Life Sciences, Pietermaritzburg Campus, University of Kwa-Zulu Natal, Pietermaritzburg, Scottsville 3209, South Africa; Department of Biotechnology and Food Technology, Faculty of Science, University of Johannesburg, Doornfontein Campus, Johannesburg 2050, South Africa

## Abstract

Over the years, the increase in antibiotic resistant *Klebsiella pneumoniae* has highlighted the urgent need for the identification of novel therapeutic compounds and targets. ClpK is a caseinolytic ATPase that plays an important role in protein quality control and thermotolerance, making it a promising drug target. This study used *in silico* methods including ADMET screening, molecular docking and molecular dynamics simulations to evaluate binding efficiency and dynamic stability of selected natural and synthetic compounds with ClpK under varying ionic conditions. ADMET screening of synthetic and natural compounds excluded compounds violating Lipinski’s rule of 5 or showing predicted toxicity. Sclerotiamide P was identified as the top binder (-10.9 kcal/mol), while compound D3 had the weakest binding (-4.5 kcal/mol). The top ten compounds with the most favourable binding scores were then further investigated using molecular dynamics simulations in the presence of NaCl and MgCl_2_. Trajectory and MM-GBSA binding analyses confirmed that ClpK-ligand complexes remained stable in both ionic environments, with negative binding free energy values indicative of favourable interactions. Overall, the study highlighted ADEP3 and sclerotiamide derivatives as promising ClpK inhibitors and provides a foundation for further experimental validation.

## 1.1. Introduction

The Gram-negative pathogen, *Klebsiella pneumoniae* has drawn the attention of many researchers following its classification as a critical priority by the World Health Organization (WHO) (Prestinaci *et al*., 2015). This opportunistic pathogen accounts for 30% of Gram-negative nosocomial infections with a mortality rate of more than 50% (Podschun and Ullmann, 1998). It can infect wounds, soft tissues, the respiratory tract and urinary tract of newborns, elderly and immunocompromised patients (Bagley, 1985, Li *et al*., 2023). *K. pneumoniae* has displayed resistance to major classes of antibiotics, including carbapenems, cephalosporins, aminoglycosides, and fosfomycin (Li *et al*., 2023, Daya *et al*., 2022). This increasing antibiotic resistance is attributed to various mechanisms including enzymatic antibiotic modification or inactivation, biofilm formation and expression of antibiotic specific efflux pumps (Li *et al*., 2023, Mulani *et al*., 2019).

The growing threat of antibiotic resistance underscores the urgent need to develop alternative therapeutic strategies to combat *K. pneumoniae* (Mulani *et al*., 2019). One such approach involves the design of inhibitors that selectively target proteins essential for its survival. Among these key survival proteins, are Caseinolytic (Clp) ATPases, which use the energy from ATP hydrolysis to drive the refolding or degradation of misfolded and aggregated proteins, thus maintaining protein quality control (Motiwala *et al*., 2022, Culp and Wright, 2017, Carroni *et al*., 2017, Lairson *et al*., 2008). Inhibiting Clp ATPases compromises protein homeostasis by increasing the level of misfolded or aggregated proteins, ultimately resulting in cell toxicity and death (Gersch *et al*., 2015). Targeting these proteins therefore offers an opportunity to disrupt critical pathways without affecting host systems.

Clp ATPases share the canonical Walker A and B motifs, which are essential for ATP binding and hydrolysis. The Walker A motif contains the GXXXXGK[T/S] consensus sequence; mutations in this region prevent nucleotide binding and lead to protein inactivation (Seraphim and Houry, 2020, Bouchnak and Van Wijk, 2021). The Walker B motif includes a conserved hydrophobic hhhhDE sequence, and mutations to this motif result in substrate trapping, where proteins bind to the chaperone but cannot be released resulting in proteotoxicity and cell death (Bouchnak and Van Wijk, 2021, Rei Liao and Van Wijk, 2019). The strict conservation of these motifs across bacterial species highlights their important role in Clp ATPase function and is consequently attractive for rational inhibitor design. Small-molecule inhibitors that disrupt ATP binding or hydrolysis at these sites could selectively inactivate Clp ATPases and protein homeostasis, providing a promising therapeutic target (Berg *et al*., 2007).

*In silico* protein-ligand docking is an effective and valuable technique to explore how candidate ligands might interact with proteins (Mfeka *et al*., 2025). By simulating binding events, docking allows for the evaluation of both the preferred orientation of ligands within the active or binding site and their relative binding affinities (Afriza *et al*., 2018, Shamsara, 2018). This approach, compared to experimental methods, is cost-effective and time-efficient, allowing for high-throughput screening of multiple ligands against target proteins (Afriza *et al*., 2018). To date, several natural and synthetic compounds targeting essential proteins in clinically relevant pathogens have been developed or isolated (Chaturvedi and Shrivastava, 2005, Plotniece *et al*., 2023, Xuan *et al*., 2023), including inhibitors of Clp ATPases (such as ClpC, and ClpX) and Clp proteases (ClpP) (Table 1).

**Table 1:** Mode of action of the compounds targeting Clp ATPases and ClpP.

| Compound | Mode of Action | Reference |
| --- | --- | --- |
| **Dihydrothiazepine 334** | Disrupts the hexameric state of methicillin-resistant *Staphylococcus aureus* (MRSA) ClpX, inhibits ATPase activity and results in the collapse of the ClpXP proteolytic complex. | (De Campos *et al.*, 2023, Fetzer *et al.*, 2017) |
| **Thiophene 336** | Blocks ClpX ATPase activity in *S. aureus* | (Fetzer *et al.*, 2017) |
| **Synthetic peptidomimetic boronate inhibitors** | Irreversibly inhibits ClpP in methicillin-resistant *S. aureus*. | (Bhandari *et al.*, 2018) |
| **Phenyl ester Compounds** | Inhibits ClpP activity via covalent modification of the active site. | (Ishikawa *et al.*, 2024, Bhandari *et al.*, 2018, Hackl *et al.*, 2015) |
| **Armeniaspirols** | Competitive inhibition of ClpXP and ClpYQ in *Bacillus subtilis* and disrupts the regulation of key proteins such as FtsZ, DivIVA, and MreB. | (Labana *et al.*, 2021) |
| **Acyldepsipeptides** | Displace ClpX ATPase by binding to the specific IGF and IGL motifs that are critical for ClpP and Clp ATPase interaction. | (Moreno-Cinos *et al.*, 2019, Kirstein *et al.*, 2009) |
| **Hydantoins** | Hydantoin 1 and 286 bind to the fully assembled ClpXP complex and inhibits protease activity. | (Fetzer *et al.*, 2019) |
| **Rufomycin B** | Inhibit *M. tuberculosis* ClpC1 and results in the modulation of protein degradation of intracellular proteins. | (Choules *et al.*, 2019) |
| **Lassomycin** | Binds to a highly acidic region of the ClpC1 ATPase complex of *M. tuberculosis* and stimulates ATPase activity without stimulating protein degradation. | (Gavrish *et al.*, 2014) |
| **Cyclomarin A** | Binds to the regulatory subunit of the *Mycobacterium* Clp protease complex resulting in elevated proteolysis and cell death. | (Schmitt *et al.*, 2011) |
| **Ecumicin** | Inhibits ClpC1 proteolytic activity whilst simultaneously stimulating ATPase activity. | (Choules *et al.*, 2019) |
| **Sclerotiamides** | Non-peptide-based natural product activator of ClpP resulting in excessive protein degradation. | (Lavey *et al.*, 2016) |

To this date, inhibitors specifically targeting ClpK remain largely unexplored. ClpK is a thermotolerant Clp ATPase, first identified in *K. pneumoniae,* with later studies confirming its presence across multiple *Klebsiella* species (Bojer *et al*., 2010, Bojer *et al*., 2012, Motiwala *et al*., 2021). To address this knowledge gap, previously identified compounds and plant derived flavonoids such as quercetin, (-)-Epigallocatechin-3-Gallate (EGCG), and kaempferol which have been identified to inhibit other Clp ATPases, heat shock proteins and mitochondrial ATPases (Khan *et al*., 2013, Zininga *et al*., 2017, Zanini *et al*., 2007) were investigated. Candidate compounds were subjected to ADMET (absorption, distribution, metabolism, excretion, and toxicity) analysis to assess pharmacokinetic and safety profiles. The filtered compounds were then docked to ClpK, and the resulting interactions were evaluated by free-binding energy calculations, molecular dynamics simulations and post-dynamic analyses.

### 1.2. Results

### 1.2.1. ClpK modelling and binding pocket identification

A homology model of ClpK (Figure 1A) was generated using the AlphaFold server. The model was evaluated for structural quality using structural parameters (Figure 1B) and displayed favourable stereochemical properties with 98.53% of residues in the most favoured region of the Ramachandran plot. Remaining parametric scores confirmed that the predicted ClpK structure was structurally reliable and suitable for further *in silico* analysis.

**Figure 1:**
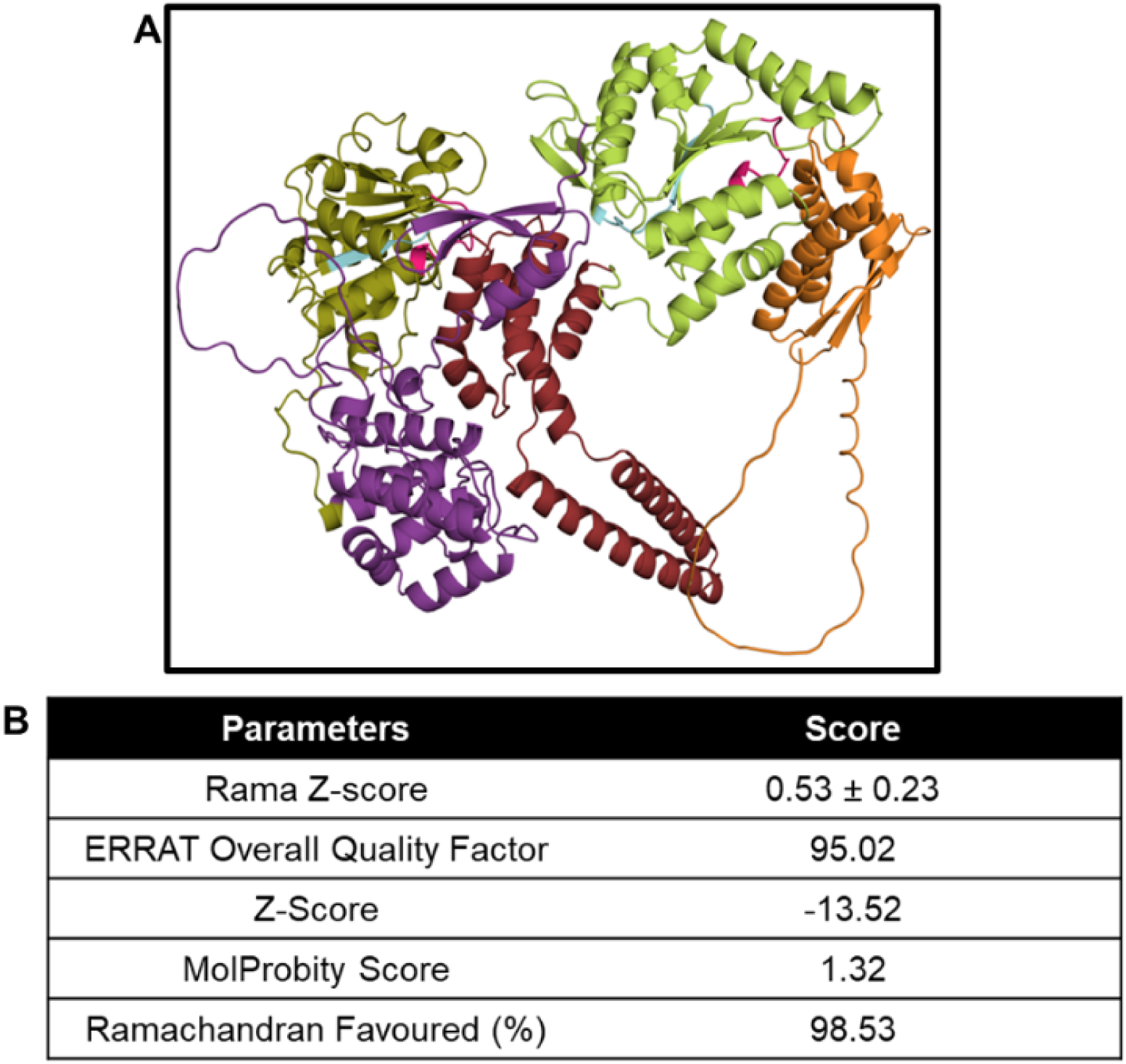
ClpK modelled using the AlphaFold server. The structure was visualised and coloured using PyMol (Schrödinger, 2015). Five domains: N-terminal domain (residues 1–244; purple); Nucleotide Binding Domain (NBD) 1 (residues 245–445; olive); the D1-small domain (residues 446–590; maroon); NBD2 (residues 591–814; lime); and the D2-small domain/ C-terminal domain (residues 815–952; orange). The Walker A and Walker B motifs were coloured hot pink and cyan, respectively.

To assess ClpK druggability prior to ligand screening, potential binding pockets were predicted using the DoGSiteScorer module on the ProteinPlus server, which identifies cavities based on a difference of Gaussian filter. A total of 29 potential binding sites were identified, of which 14 binding sites had a druggability score greater than 0.5. Among these 5 models exhibited a SimpleScore greater than 0.5, indicating favourable size, shape, and flexibility for ligand binding (Figure 2A, 2B) (Volkamer *et al*., 2010). These pockets contained hydrophobic regions that facilitate protein-protein or protein-molecule interactions (Jernigan *et al*., 2022), as well as hydrogen acceptors and donors that further facilitate ligand binding (Zhao and Huang, 2011) (Figure 2A). Collectively, these features confirmed ClpK as a druggable target suitable for subsequent docking analyses.

**Figure 2:**
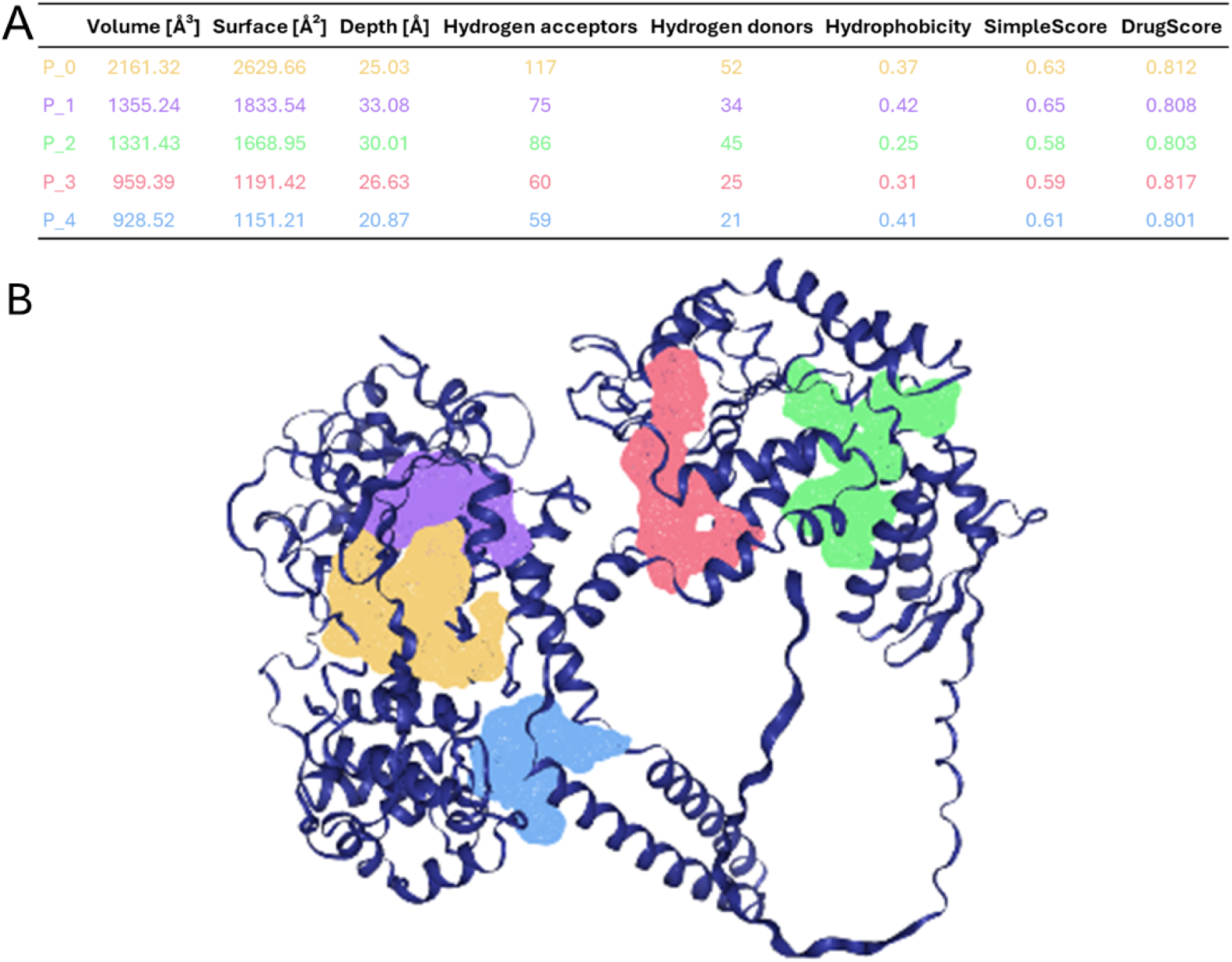
Prediction of the potential binding pockets in ClpK. A) Quantitative properties of the predicted binding pockets. B) Visual representation of the top five predicted binding pockets on the ClpK. Each pocket is colour coded to correspond with the data in panel A: P_0 (yellow), P_1 (purple), P_2 (green), P_3 (red), and P_4 (blue). The ClpK backbone is shown in a cartoon representation as navy blue.

### 1.2.2. ADMET analysis

To assess oral bioavailability and drug likeliness, the compounds were analysed according to Lipinski’s ‘rule of five,’ which outlines key physicochemical criteria for drug-like behaviour (Varma *et al*., 2006). According to these rules, a compound is considered drug-like if it meets at least two of the following four criteria; i) molecular weight less than 500, ii) calculated LogP (cLogP) value less than 5, iii) less than 10 H-bond acceptors and iv) less than 5 H-bond donors (Ali *et al*., 2023). The cLogP value indicates the hydrophilic (negative values) or lipophilic nature of a compound (Silakari and Singh, 2021). All selected small ligands satisfied these criteria, indicating potentially suitability for oral administration (Table 2).

**Table 2:** ADME properties of small ligands predicted using the SWISSADME server.

| Compound | Molecular Weight (g/mol) | cLogP | H-bond Donor | H-bond Acceptor | Lipinski's Rule |
| --- | --- | --- | --- | --- | --- |
| 334 | 357.43 | 3.87 | 2 | 2 | 4 |
| 336 | 347.45 | 4.29 | 1 | 1 | 4 |
| G2 | 180.24 | 2.59 | 0 | 2 | 4 |
| U1 | 300.44 | 5.18 | 0 | 2 | 3 |
| AV126 | 374.41 | 3.56 | 0 | 6 | 4 |
| AV145 | 353.44 | 2.89 | 1 | 5 | 4 |
| AV170 | 374.38 | 3.26 | 0 | 7 | 4 |
| D3 | 248.36 | 4.21 | 0 | 2 | 4 |
| E2 | 230.26 | 2.56 | 0 | 3 | 4 |
| **Armeniaspirols** |  |  |  |  |  |
| Armeniaspirol-A | 384.25 | 3.72 | 1 | 4 | 4 |
| Armeniaspirol -B | 384.25 | 3.59 | 1 | 4 | 4 |
| Armeniaspirol -C | 398.28 | 3.89 | 1 | 4 | 4 |
| 5-chloro- armeniaspirols | 418.70 | 4.26 | 1 | 4 | 4 |
| **Acyldepsipeptide antibiotics (ADEPs)** |  |  |  |  |  |
| ADEP1 | 718.84 | 1.73 | 3 | 8 | 3 |
| ADEP2 | 798.92 | 3.10 | 3 | 10 | 3 |
| ADEP3 | 798.92 | 2.90 | 3 | 10 | 3 |
| ADEP4 | 770.86 | 2.75 | 3 | 10 | 3 |
| ADEP5 | 843.91 | 1.71 | 4 | 13 | 2 |
| IDR-10001 | 744.83 | 2.49 | 3 | 10 | 3 |
| IDR-10011 | 790.92 | 2.64 | 3 | 10 | 3 |
| **Hydantoin** |  |  |  |  |  |
| Mephenytoin | 218.25 | 1.48 | 1 | 2 | 4 |
| (+)-Hydantocidin | 218.16 | -2.33 | 5 | 6 | 4 |
| Azimilide | 457.95 | 2.71 | 0 | 6 | 4 |
| Dantrolene | 314.25 | 0.94 | 1 | 6 | 4 |
| Fosphenytoin | 362.27 | 0.69 | 3 | 6 | 4 |
| Hydantoin-1 | 481.99 | 4.13 | 1 | 3 | 4 |
| Hydantoin-286 | 432.49 | 3.42 | 3 | 4 | 4 |
| Iprodion | 330.17 | 2.42 | 1 | 3 | 4 |
| Nifurtoinol | 268.18 | -0.83 | 1 | 7 | 4 |
| Nilutamide | 317.22 | 1.66 | 1 | 7 | 4 |
| Nitrofurantoin | 238.16 | -0.50 | 1 | 6 | 4 |
| Phenytoin | 252.27 | 1.81 | 2 | 2 | 4 |
| Sorbinil | 236.20 | 0.99 | 2 | 4 | 4 |
| **Flavonoids** |  |  |  |  |  |
| Quercetin | 302.24 | 1.23 | 5 | 7 | 4 |
| Epigallocatechin gallate (EGCG) | 458.37 | 0.95 | 8 | 11 | 2 |
| Kaempferol | 286.24 | 1.58 | 4 | 6 | 4 |
| Fisetin | 286.24 | 1.55 | 4 | 6 | 4 |
| Apigenin | 270.24 | 2.11 | 3 | 5 | 4 |
| Luteolin | 286.24 | 1.73 | 4 | 6 | 4 |
| Genistein | 270.24 | 2.04 | 3 | 5 | 4 |

The selected small ligands demonstrated distinct pharmacokinetic and toxicity profiles, as determined by SWISSADME and ProTox 3.0 analyses. Variations were observed in the predicted bioavailability, absorption, synthetic accessibility, BBB permeability, and toxicity indicating key differences in druggability and safety. The bio-availability score estimates how much of an orally administered compound reaches the blood; a value closer to 1 indicates better bioavailability (Martin, 2005). Among the test ligands, acyldepsipeptide antibiotics (ADEPs) and epigallocatechin gallate (EGCG) had a bioavailability score of 0.17, indicating poor bioavailability, whereas the other ligands had scores >0.5, suggesting good bioavailability (Daina *et al*., 2017). Gastrointestinal (GI) absorption was high, except for ADEPs, EGCGs and (+)- hydantocidin. Synthetic accessibility scores were less than 5 for all except ADEPs, suggesting ease of synthesis. BBB permeability was observed for nine ligands, including G2 and D3. Toxicity classification ranged from Class 1 (most toxic, fatal if swallowed, ≤ 5 mg/kg) to Class 6 (non-toxic, > 5000 mg/kg); armeniaspirols A - C were predicted to be highly toxic and were excluded from further analysis (Table 3) (Ali *et al*., 2023, Alazmi and Motwalli, 2021).

**Table 3:** ADMET properties of small ligands predicted using the SWISSADME and ProTox 3.0 server.

| Compound | Bioavailability score | Gastrointestinal (GI) absorption | Synthetic accessibility | Blood brain barrier (BBB) permeability | LD50 (mg/kg) | Toxicity class |
| --- | --- | --- | --- | --- | --- | --- |
| 334 | 0.55 | High | 4.19 | No | 1000 | 4 |
| 336 | 0.55 | High | 4.21 | No | 1000 | 4 |
| G2 | 0.55 | High | 3.29 | Yes | 8000 | 6 |
| U1 | 0.55 | High | 3.26 | Yes | 2100 | 5 |
| AV126 | 0.55 | High | 3.05 | No | 4500 | 5 |
| AV145 | 0.55 | High | 3.30 | No | 500 | 4 |
| AV170 | 0.55 | High | 2.75 | No | 1000 | 4 |
| D3 | 0.55 | High | 3.71 | Yes | 5000 | 5 |
| E2 | 0.55 | High | 3.05 | Yes | 2000 | 4 |
| **Armeniaspirols** |  |  |  |  |  |  |
| Armeniaspirol-A | 0.55 | High | 4.49 | Yes | 4 | 1 |
| Armeniaspirol -B | 0.55 | High | 4.75 | Yes | 4 | 1 |
| Armeniaspirol -C | 0.55 | High | 4.86 | Yes | 4 | 1 |
| 5-chloro-armeniaspirols | 0.55 | High | 4.48 | Yes | 360 | 4 |
| **Acyldepsipeptide antibiotics (ADEPs)** |  |  |  |  |  |  |
| ADEP1 | 0.17 | Low | 7.13 | No | 200 | 3 |
| ADEP2 | 0.17 | Low | 7.40 | No | 200 | 3 |
| ADEP3 | 0.17 | Low | 7.40 | No | 200 | 3 |
| ADEP4 | 0.17 | Low | 7.31 | No | 200 | 3 |
| ADEP5 | 0.17 | Low | 7.67 | No | 200 | 3 |
| IDR-10001 | 0.17 | Low | 7.14 | No | 200 | 3 |
| IDR-10011 | 0.17 | Low | 7.27 | No | 200 | 3 |
| **Hydantoins** |  |  |  |  |  |  |
| Mephentoin | 0.55 | High | 2.19 | Yes | 440 | 4 |
| (+)-Hydantocidin | 0.55 | Low | 4.16 | No | 2000 | 4 |
| Azimilide | 0.55 | High | 4.18 | Yes | 1000 | 4 |
| Dantrolene | 0.55 | High | 3.27 | No | 2300 | 5 |
| Fosphenytoin | 0.56 | High | 2.92 | No | 500 | 4 |
| Hydantoin-1 | 0.55 | High | 3.91 | No | 1300 | 4 |
| Hydantoin-286 | 0.55 | High | 3.52 | No | 200 | 3 |
| Iprodion | 0.55 | High | 2.57 | Yes | 3500 | 5 |
| Nifurtinol | 0.55 | High | 3.19 | No | 1000 | 4 |
| Nilutamide | 0.55 | High | 2.33 | No | 625 | 4 |
| Nitrofurantoin | 0.55 | High | 3.10 | No | 1000 | 4 |
| Phenytoin | 0.55 | High | 1.83 | Yes | 150 | 3 |
| Sorbinil | 0.55 | High | 3.27 | No | 5000 | 5 |
| **Flavonoids** |  |  |  |  |  |  |
| Quercetin | 0.55 | High | 3.23 | No | 159 | 3 |
| Epigallocatechin gallate (EGCG) | 0.17 | Low | 4.20 | No | 1000 | 4 |
| Kaempferol | 0.55 | High | 3.14 | No | 3919 | 5 |
| Fisetin | 0.55 | High | 3.16 | No | 159 | 3 |
| Apigenin | 0.55 | High | 2.96 | No | 2500 | 5 |
| Luteolin | 0.55 | High | 3.02 | No | 3919 | 5 |
| Genistein | 0.55 | High | 2.87 | No | 2500 | 5 |

Among the large ligands, only Sclerotiamide derivatives fully complied with Lipinski’s rule of five, while other ligands violated one or more criteria. Maximum tolerated doses were less than 5 mg/kg/day, indicating high oral toxicity and suggesting the need for non-oral administration (Ali *et al*., 2023). Most ligands were lipophilic (logP > 0), supporting their formulation for non-oral routes, such as injections (Han *et al*., 2021). Sclerotiamides demonstrated logBB ≥ -1, while others showed poor penetration (logBB < -1). Sclerotiamide L was predicted to be toxic and excluded from subsequent analysis (Table 4).

**Table 4:**
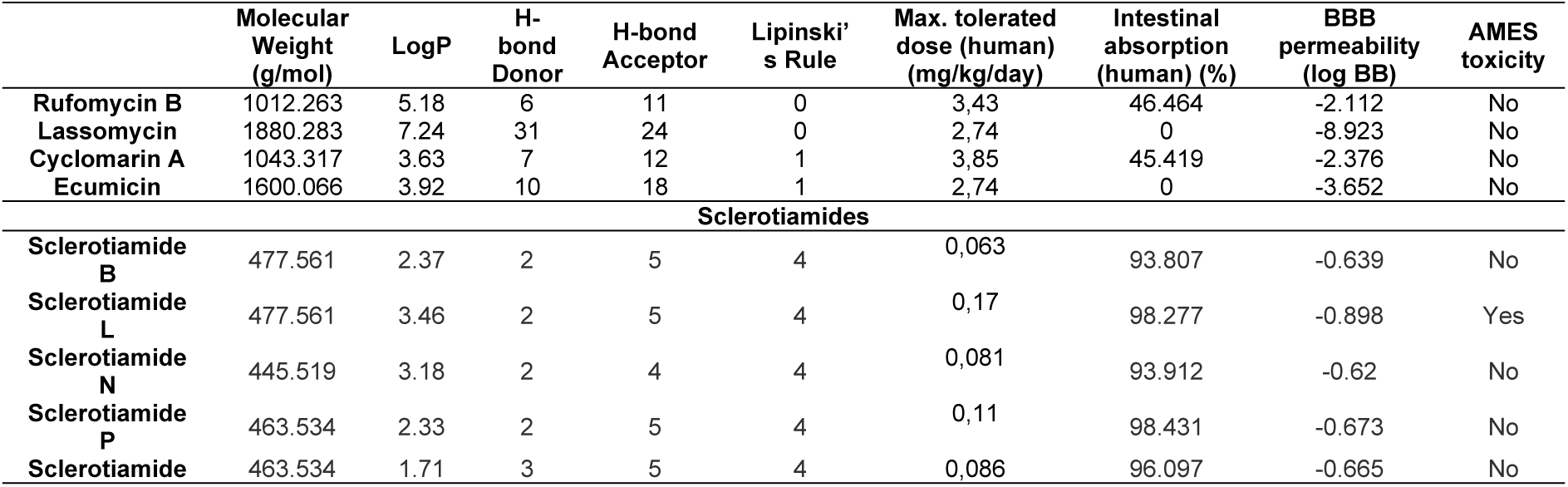
ADMET analysis of the large ligands predicted using the pkCSM server.

### 1.2.3. Molecular Docking studies

Molecular docking analysis revealed binding affinities ranging from -10.9 kcal/mol (Sclerotiamide P) to -4.5 kcal/mol (compound D3) indicating varying binding strengths (Table 5). The top ten compounds with the most favourable (negative) binding affinities were selected for subsequent *in silico* analysis, including molecular dynamics simulation and binding free energy calculations.

**Table 5:** Molecular docking scores for ClpK-ligand complexes as obtained using AutoDock Vina on the PyRx software.

| Compound | Binding affinity (kcal/mol) | Compound | Binding affinity (kcal/mol) |
| --- | --- | --- | --- |
| Sclerotiamide P | -10.9 | Luteolin | -7.7 |
| Sclerotiamide | -10.5 | Rufomycin B | -7.6 |
| ADEP3 | -10.2 | ADEP4 | -7.5 |
| Sclerotiamide N | -10.1 | Azimilide | -7.5 |
| ADEP1 | -9.9 | Genistein | -7.5 |
| Hydantoin-1 | -9.1 | ADEP4 | -7.5 |
| Sclerotiamide B | -9.0 | AV126 | -7.4 |
| Epigallocatechin gallate (EGCG) | -9.0 | Iprodion | -7.4 |
| ADEP5 | -8.9 | Cyclomarin A | -7.3 |
| ADEP2 | -8.5 | 5-chloro- armeniaspirols | -7.0 |
| IDR-10011 | -8.4 | Nifurtinol | -7.0 |
| Hydantoin-286 | -8.4 | AV145 | -6.9 |
| Fisetin | -8.4 | Nitrofurantoin | -6.7 |
| 336 | -8.2 | AV170 | -6.5 |
| IDR-10001 | -8.2 | Mephentyoin | -6.5 |
| Fosphenytoin | -8.2 | Lassomycin | -6.5 |
| Phenytoin | -8.1 | Ecumicin | -6.5 |
| Kaempferol | -8.1 | Sorbinil | -6.5 |
| 334 | -7.9 | (+)-Hydantocidin | -6.4 |
| Quercetin | -7.9 | U1 | -6.2 |
| Dantrolene | -7.7 | E2 | -5.5 |
| Nilutamide | -7.7 | G2 | -5.0 |
| Apigenin | -7.7 | D3 | -4.5 |

The binding interactions of the top 10 ligands and ClpK were visualised in Figure 3. All ligands formed multiple stabilising contacts with ClpK, including hydrogen bonds, π-π stacking, amide-π, and hydrophobic interactions, consistent with their docking affinities. Sclerotiamides and ADEP5 predominantly bind within Pocket 0 (N-terminal domain, NBD1, and D1-small domain), EGCG and ADEP1 bind within Pocket 3 (D1-small domain and NDB2 region), hydantoin and ADEP3 interacts in Pocket 1 (primarily engaging NBD1), and ADEP2 in Pocket 4 (making contacts within the N-terminal domain) (Figure 2, Figure 3). The following residues participated in multiple ligand interactions: PRO283, THR462, ASP282, LEU108, PHE197, VAL198, and GLY279, each involved in five or more ligands, suggesting their role as core binding sites thus suggesting their potential role in ligand recognition and stabilisation.

**Figure 3:**
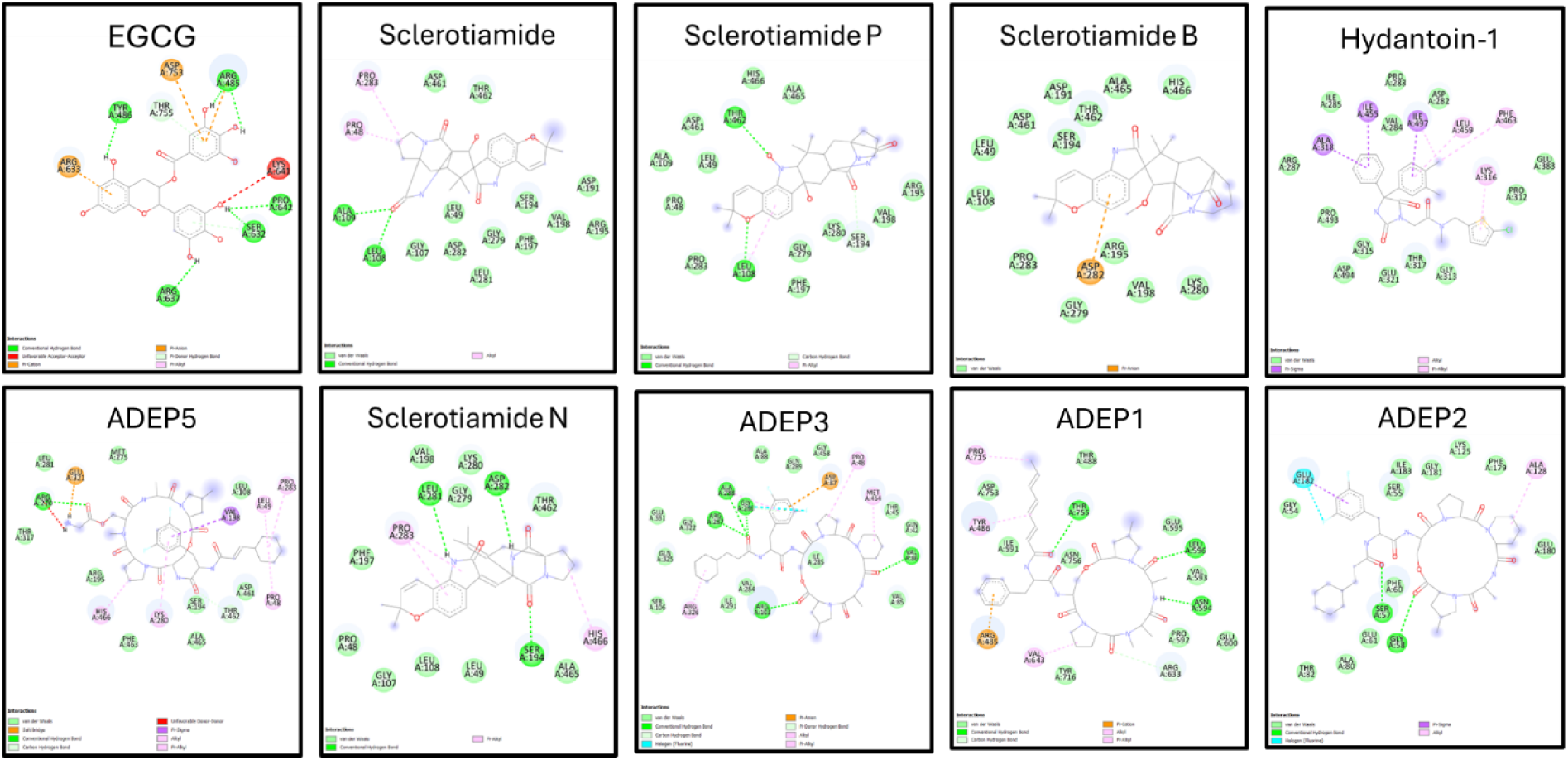
Molecular interaction of ClpK with the top 10 ligands exhibiting the strongest binding affinities. Hydrogen bonds are represented by green balls and sticks, π-π/π-sigma/amide-π interactions are represented by violet balls and sticks, π-alkyl/alkyl interactions are presented by pink balls and sticks, π-sulphur bonds are represented by gold balls and sticks, and carbon-hydrogen bonds are represented as light green balls and sticks.

### 1.2.4. Molecular dynamics simulations

MD simulations of 100 ns were performed to investigate ClpK-ligand interactions, stability, and conformational changes. The potential energy of a system indicates the energetic stability of the *apo*- and complexed ClpK structures (Ajao *et al*., 2012). The average potential energy of apo-ClpK was observed to be -2813010.92 kJ/mol in the presence of NaCl, and this increased slightly to -2859433.80 kJ/mol in the presence of MgCl_2,_ indicating a stable and well-equilibrated system under both ionic conditions. The increase in potential energy observed in the presence of compounds is expected as ligand binding would result in an increase in the systems total energy (Figure 4). Despite these differences, the potential energy remains stable over the 100 ns simulation period suggesting that the systems reached equilibration and maintained energetic stability.

**Figure 4:**
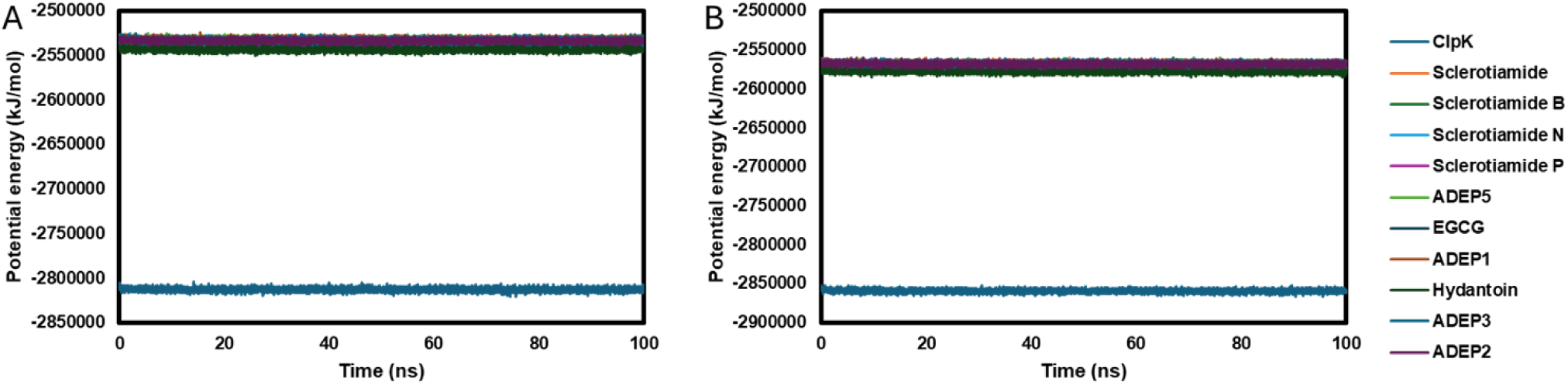
Potential energy (kJ/mol) profiles of *apo*-ClpK and ClpK-compound complexes analysed over 100 ns. In the presence of **A)** NaCl and **B)** MgCl_2_.

RMSD analysis showed an average of 0.941 nm for *apo*-ClpK in NaCl, increasing to 1.21 nm in MgCl_2_, indicating greater structural deviations and enhanced flexibility in the presence MgCl_2_ (Figure 5). Ligand binding increased RMSD in NaCl (1.01 nm to 1.62 nm), with ClpK-Sclerotiamide N showing the highest deviation (1.62 nm). In MgCl_2_, some complexes, notably ClpK-Sclerotiamide-B (1.14 nm) and ClpK-ADEP3 (1.06 nm), exhibited decreased RMSD, indicating stable ligand binding, while others, such as ClpK-ADEP2, displayed increased RMSD up to 1.76 nm, suggesting destabilisation (Figure 5).

**Figure 5:**
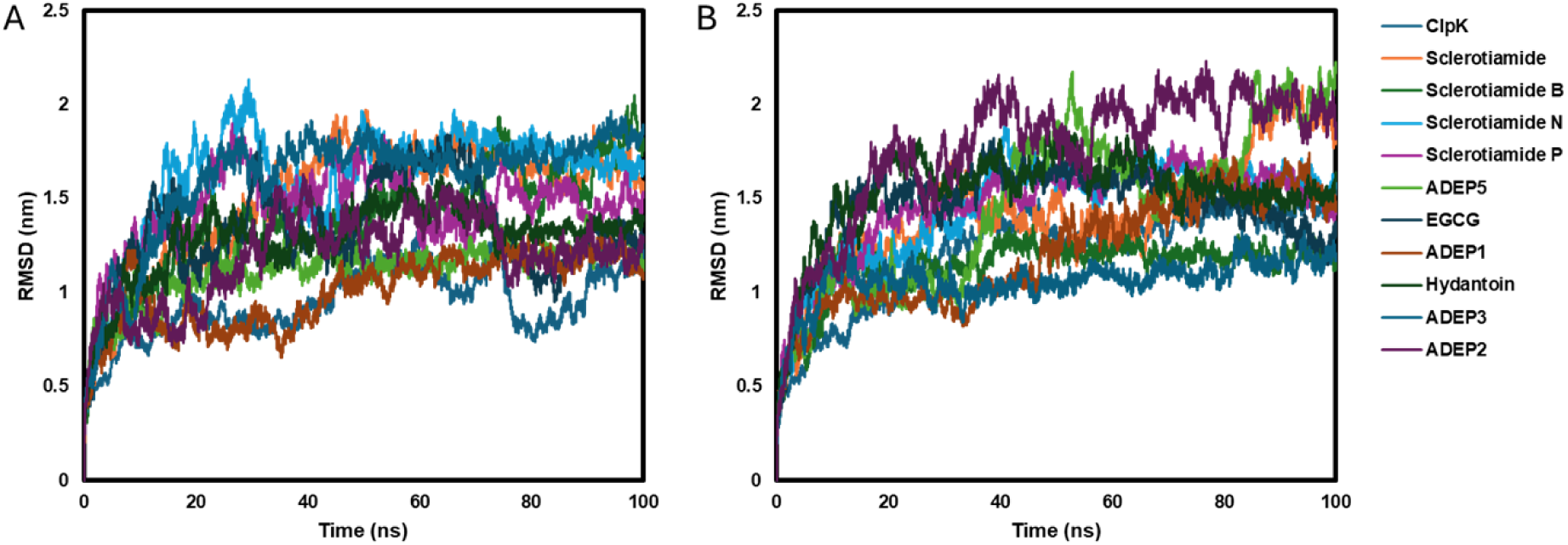
RMSD profiles of *apo*-ClpK and ClpK-compound complexes over 100 ns molecular dynamics simulations. Simulations were run in the presence of **A)** NaCl and **B)** MgCl_2_.

Residue flexibility over the 100 ns simulation was assessed using RMSF (Figure 6). For apo-ClpK, average RMSF increased slightly from 0.37 nm in NaCl to 0.38 nm in MgCl_2_, consistent with the RMSD results, indicating that MgCl_2_ affects structural stability. Ligand binding increased RMSF, ranging from 0.42 nm to 0.64 nm in NaCl and 0.41 nm to 0.62 nm in MgCl_2_. In NaCl, fluctuations were distributed across N-terminal, middle, and C-terminal domains, with Sclerotiamide N exhibiting the highest RMSF (0.64 nm), reflecting destabilisation. In MgCl_2,_ fluctuations were primarily at the C-terminal, with ADEP5 (0.62 nm) and ADEP2 (0.61 nm) complexes showing higher RMSF, corresponding to the destabilising effect observed in RMSD analysis.

**Figure 6:**
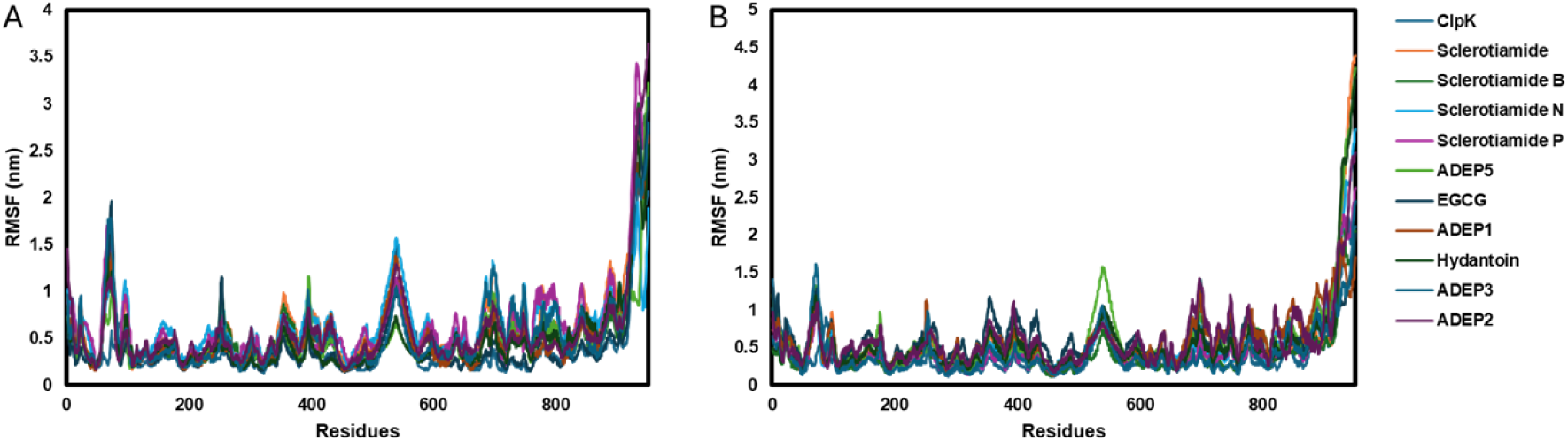
RMSF profiles of *apo*-ClpK and ClpK-compound complexes analysed over 100 ns. In the presence of **A)** NaCl and **B)** MgCl_2_.

The Rg was used to assess the structural compactness of ClpK (Figure 7). A*po*-ClpK maintained stable Rg values of 3.83 nm in NaCl and 3.85 nm in MgCl_2_. In NaCl, ligands such as Sclerotiamide B, ADEP1, ADEP5, and Hydantoin reduced Rg, indicating tighter conformations. In contrast, Sclerotiamide, Sclerotiamide N, Sclerotiamide P, EGCG, ADEP3 and ADEP2 showed increased Rg, suggesting loosening or partial unfolding. In MgCl_2_, most ligands maintained Rg values comparable to apo. However, Sclerotiamide B, ADEP1 and ADEP3 showed decreased Rg, while Sclerotiamide P, Hydantoin and ADEP2, suggesting Mg^2+^-dependent differences in ligand-induced relaxation.

**Figure 7:**
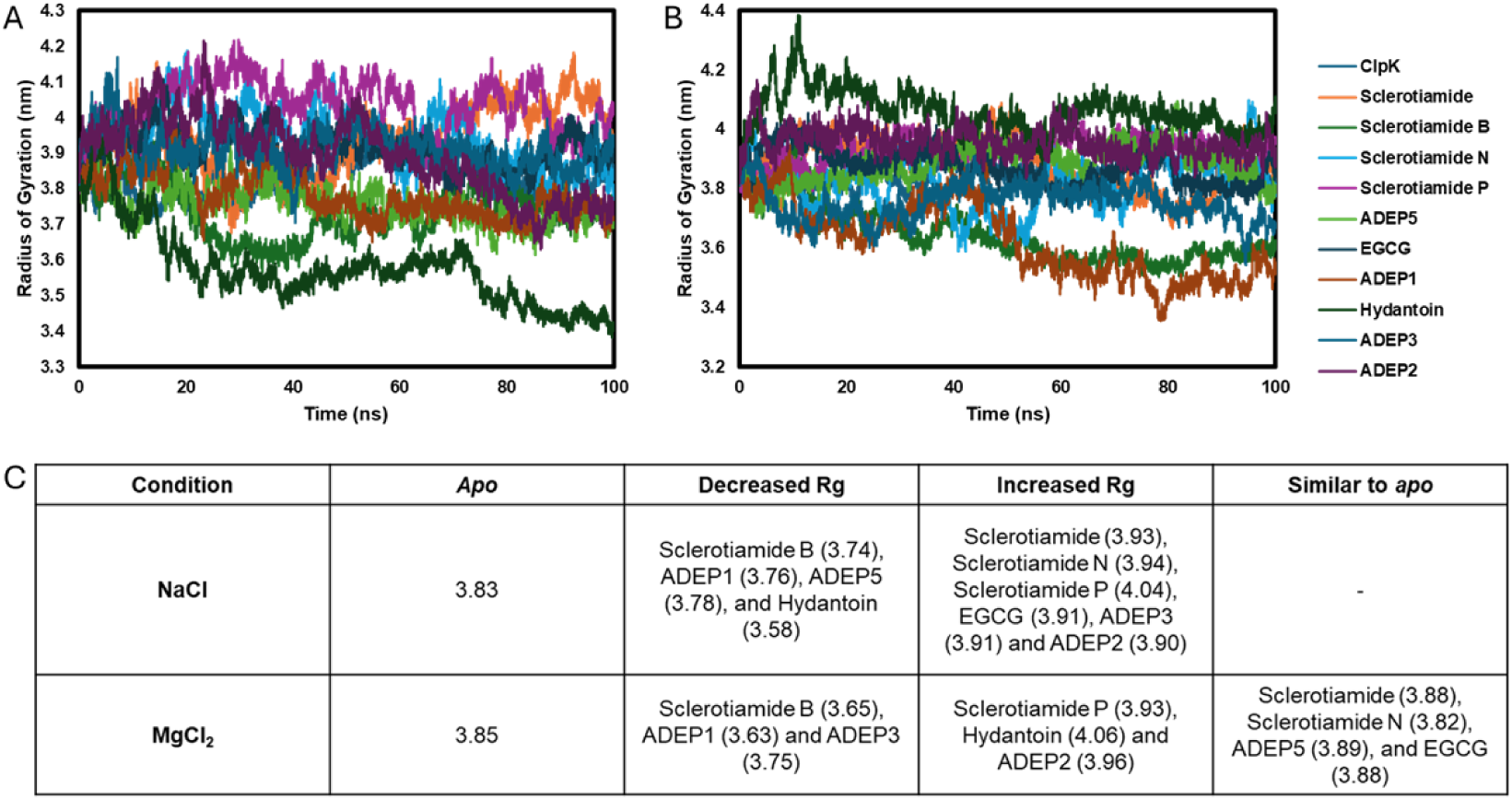
Radius of gyration (Rg) profiles of *apo*-ClpK and ClpK-compound complexes analysed over 100 ns. In the presence of **A)** NaCl and **B)** MgCl_2_. **C)** Mean Rg values (nm) of ClpK in *apo* and ligand bound states.

The SASA represents the portion of the protein that is exposed to surrounding solvent molecules and is important for protein solvation and compactness analysis (Borjian Boroujeni *et al*., 2021). The average SASA of apo-ClpK was observed to be 514.25 nm/S^2^/N in the presence of NaCl and 504.86 nm/S^2^/N in the presence of MgCl_2_, indicating a slightly more compact conformation under MgCl_2_ conditions. A lower SASA indicates reduced solvent exposure, this is associated with tighter protein packing and correlates with the slight increase in Rg observed in Figure 6 in the presence of MgCl_2_. In the presence of ligands, an increase in average SASA was observed under both ionic conditions. This increase could be attributed to ligand induced conformational changes in the surface-exposed regions of ligand binding sites which may cause protein unfolding or expansion of the surface area (Figure 8). Additionally, increased SASA values indicating structural relaxation and therefore a decrease in protein stability (Borjian Boroujeni *et al*., 2021).

**Figure 8:**
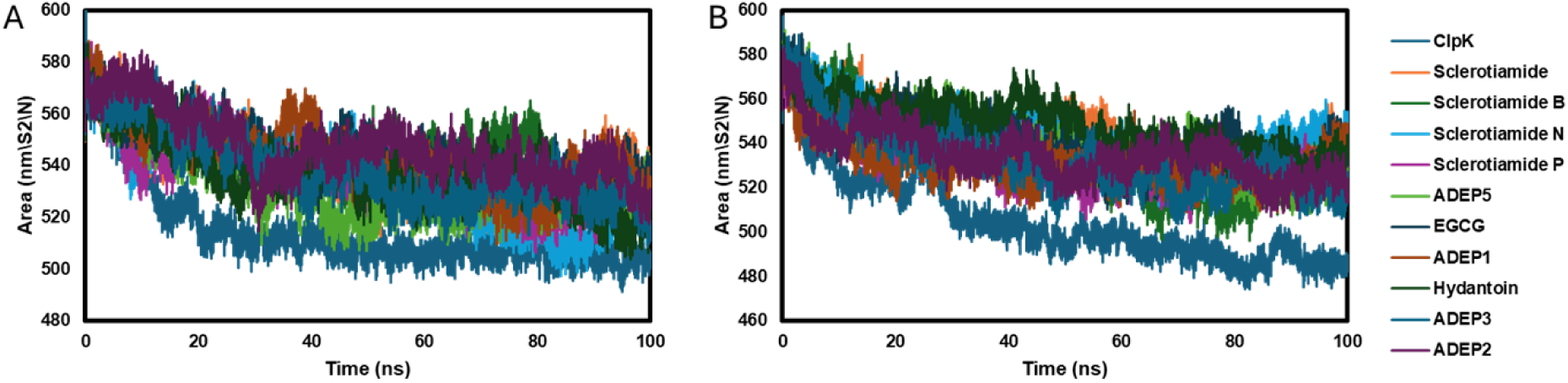
SASA profiles of *apo*-ClpK and ClpK-compound complexes analysed over 100 ns. In the presence of **A)** NaCl and **B)** MgCl_2_.

The number of hydrogen bonds between proteins and ligands can be investigated to evaluate interaction stability (Molaakbari *et al*., 2024). To evaluate the dynamic nature of these interactions, the number and persistence was investigated in the presence of NaCl and MgCl_2_ (Figure 9). A high diversity and fluctuation of hydrogen bonds is observed in the presence of NaCl (Figure 9A), however in the presence of MgCl_2_ the hydrogen bonds are more stable and persistent (Figure 9B). The MD simulations indicate that the ionic environment and ligand binding influence the global structural dynamics of ClpK.

**Figure 9:**
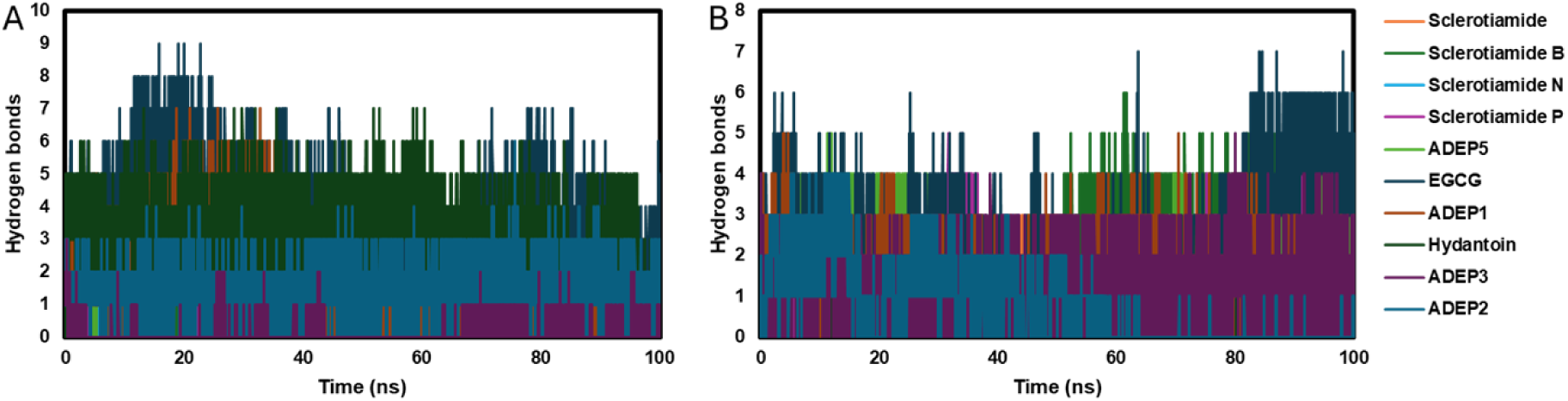
Hydrogen bond count between ClpK and compounds over the course of molecular dynamic simulations. The number of hydrogen bonds formed was investigated in the presence of **A)** NaCl and **B)** MgCl_2_.

Table 6 summarises the binding free energies values for the ClpK-compound complexes in the presence of NaCl and MgCl_2_, as calculated using MM-GBSA. Negative values indicate favourable binding. In the presence of NaCl, ADEP3 (-41.39 kcal/mol) had the lowest binding energy, followed by Sclerotiamide (-35.65 kcal/mol) and ADEP1 (-35.43 kcal/mol). Similarly, in the presence of MgCl_2_, ADEP3 (-42.59 kcal/mol) remained the most favourable binder, followed by ADEP2 (-34.03 kcal/mol) and Sclerotiamide B (27.70 kcal/mol). Interestingly, ADEP3 had the lowest binding affinity in the presence of both NaCl and MgCl_2_ suggesting that ADEP3 may be the most stable and effective binder to ClpK among the tested compounds. The binding energies of some compounds varied significantly between some compounds.

**Table 6:** Molecular Mechanics Generalized Born Surface Area (MM-GBSA) analysis of the binding free energy of ClpK and compounds.

|  | NaCl (kcal/mol) | MgCl<sub>2</sub> (kcal/mol) |
| --- | --- | --- |
| ADEP1 | -35.43 | -26.28 |
| ADEP2 | -17.26 | -34.03 |
| ADEP3 | -41.39 | -42.59 |
| ADEP5 | -22.17 | -27.13 |
| EGCG | -23.93 | -23.24 |
| Hydatoin | -31.50 | -16.07 |
| Sclerotiamide | -35.65 | -20.93 |
| Sclerotiamide B | -16.76 | -27.70 |
| Sclerotiamide N | -17.38 | -17.12 |
| Sclerotiamide P | -24.70 | -20.34 |

## 1.3. Discussion

*K. pneumoniae* is a multidrug resistant, WHO-designated priority pathogen that urgently requires the development of new therapeutic strategies. The ClpK ATPase, implicated in protein homeostasis and stress tolerance, represents a potential target for therapeutic development (Shrivastava *et al*., 2018, Abbas *et al*., 2024). In this study, *in silico* approaches were used to investigate the binding efficiency and dynamic stability of selected natural and synthetic compounds with ClpK under varying ionic conditions.

Binding site prediction in the ClpK homology model revealed 14 potential binding pockets, five of which were classified as druggable based on the pocket size and SimpleScore metrics (Figure 2). Computational binding sites prediction provides a valuable first step in drug discovery however, they may be limited by data quality and standardisation challenges (Stratiichuk *et al*., 2025). To explore the functional significance of these predicted sites, molecular docking and molecular-dynamics simulations were subsequently performed to evaluate ligand binding, interaction stability, and potential inhibitory effects. A preliminary screening of natural and synthetic compounds (Tables 2-4) enabled the exclusion of molecules with unfavourable ADMET characteristics, refining the selection for downstream docking and dynamics studies. Docking analysis identified several high affinity compounds, mainly the sclerotiamide derivates and ADEP analogues (Table 4) and this is consistent with findings that reported sclerotiamide activates ClpP, while ADEP can act as both a ClpP activator and inhibitor (Vass and Chien, 2016, Lavey *et al*., 2016).

To further characterise binding, ligand interaction profiling was conducted for the top ten compounds. Compounds were observed to form hydrogen bonds, π-π stacking, and hydrophobic interactions with residues within key functional domains of ClpK including, the N-terminal domain, NBD1, D1-small domain, and NBD2 (Figure 3). These interactions reinforced binding stability and specificity with hydrogen bonds orientating the ligands, π-π stacking enhancing affinity and hydrophobic contacts enhancing stability (Madushanka *et al*., 2023, Brylinski, 2018, Ferenczy and Kellermayer, 2022). Additionally, the involvement of multiple domains is advantageous for drug development potentially offering avenues for the development of compounds that disrupt ClpK’s interdomain interaction and ATPase function (Ferrari *et al*., 2003).

The structural dynamics of these multi-domain interactions were investigated using 100 ns MD simulations with NaCl and MgCl_2_ to explore how these ionic conditions effect structural stability and flexibility. Since ionic environments can alter protein electrostatics, conformational flexibility, and ligand stability, this comparison can provide an insight into the physiochemical factors that can influence inhibitor binding (Schönichen *et al*., 2013). The potential energy profiles of *apo*-ClpK and ligand-bound complexes remained stable throughout the simulations (Figure 4). Suggesting that complexes were energetically and thermodynamically stable, allowing for the reliable interpretation of structural and dynamic parameters (Ajao *et al*., 2012).

RMSD trajectories (Figure 5) revealed distinct stability profiles under the two ionic conditions. In the presence of NaCl, the average RMSD values increased upon ligand binding, with the sclerotiamide N complex exhibiting the highest deviation. In contrast, systems exhibited slightly higher RMSD fluctuation in MgCl_2_ suggesting enhanced overall flexibility. This indicates that the stronger electrostatic influence of Mg^2+^ ions may perturb intramolecular interactions and promote local rearrangements in the protein backbone (Schauss *et al*., 2021). This observation is consistent with findings reported by Shao *et al*. (2022), where simulations of protein 1BBL in the presence of MgCl_2_ also showed increased RMSD values, indicating enhanced protein dynamics at moderate salt concentrations. In the presence of MgCl_2_, Sclerotiamide B and ADEP3 complexes maintained stable ligand binding, while ADEP2 increased average RMSD values suggesting destabilisation. Overall, RMSD values showed that most complexes reached equilibrium within the 100 ns timeframe.

RMSF analysis (Figure 6) was consistent with the RMSD findings and provided an insight into local dynamics. A moderate increase in residue level fluctuations were observed in the presence of NaCl, with the binding of Sclerotiamide N exhibiting the highest fluctuation. In contrast, elevated peaks were observed in the presence of MgCl_2_ particularly in the C-terminal region. Increased fluctuations observed for the ADEP2 and ADEP5 complexes indicated that specific ligand interactions may induce local destabilisation in the presence of MgCl_2_. Complementary analysis of Rg (Figure 7) further investigated the influence of ions on the overall structural organisation. ClpK-ligand complexes maintained relatively consistent Rg values in the presence of NaCl, indicating preserved structural compactness and stable folding (Motiwala *et al*., 2025). In contrast, higher Rg values with more pronounced fluctuations were observed in the presence of MgCl_2_ indicating increased structural flexibility. This corresponds with the observed RMSD trends which indicate that the Mg^2+^ ions promote a more relaxed conformational state. However, all complexes reached equilibrium by the end of the 100 ns simulation time, indicating overall complex stability under both ionic conditions suggesting that ionic composition modulates ClpK-ligand interactions.

The ion-dependent difference in protein-ligand interaction stability were further investigated through SASA analysis (Figure 8). The average SASA of apo-ClpK decreased in the presence of MgCl_2_ suggesting that a more compact protein conformation is promoted under these conditions. This is also observed for the ClpK-ligand complexes where lower SASA values in the presence of MgCl_2_ suggest reduced solvent exposure and enhances structural compactness. In contrast, in the presence of NaCl higher and more fluctuating SASA values are observed therefore indicating structural relaxation (Borjian Boroujeni *et al*., 2021). This variability can further be supported through hydrogen bonding analyses (Figure 9). It was observed that NaCl promoted more diverse and fluctuating hydrogen bond patterns suggesting dynamic and transient interactions. In contrast, stable and persistent hydrogen bonds were observed in the presence of MgCl_2_ suggesting stronger electrostatic stabilisation and reduced conformational mobility. These structural observations were consistent with MM-GBSA binding free energy results (Table 5) where the negative binding free energy values of all complexes indicated that the compounds bind to ClpK under both ionic conditions, supporting their potential biological relevance. Additionally, these findings highlight the importance of carefully considering buffer composition in molecular simulations and drug screening workflows due to the influence ionic interactions can have on predicted binding behaviours.

In conclusion, this study provides a foundation for future *in vitro* studies that focus on validating the predicted interactions. To validate the *in silico* findings, it is important to assess ClpK ATPase activity in the presence of these ligands. These experiments should be performed under varying physiological conditions, including different pH levels and salt concentrations to reflect the dynamic intracellular environment. Additionally, the impact of these compounds on ClpK thermal stability would offer insights into their potential to function as stabilisers or destabilisers of ClpK. Collectively, these studies would help confirm the biological relevance of the predicted interactions and further support the development of ClpK as a promising therapeutic target against *K. pneumoniae*.

## 1.4. Materials and Methods

### 1.4.1. ClpK Modelling and Analysis

ClpK was modelled using the AlphaFold server, which predicts three-dimensional (3D) structures using the primary protein sequence (Jumper *et al*., 2021, Schwede *et al*., 2003, Ko *et al*., 2012). The modelled structure was evaluated using the MolProbity, VADAR (Volume, Area, Dihedral Angle Reporter) version 1.8, Protein Structure Analysis (ProSA) web server, ERRAT module on the saves Saves v1.6 and the SWISS MODEL structure assessment tool (Williams *et al*., 2018, Wiederstein and Sippl, 2007, Waterhouse *et al*., 2018, Willard *et al*., 2003, Colovos and Yeates, 1993). ClpK binding sites were identified using the DoGSiteScorer module on the ProteinPlus server (https://proteins.plus/, Accessed 12 August 2024).

### 1.4.2. Ligand preparation

The ligands dihydrothiazepine 334 and 336 were drawn using the ChemSketch software while the ADEP1 structure was derived from PDB ID: 3KT and sketched on the ChemSketch software using the SMILES obtained from the Protein Data Bank (PDB). Additional ligands namely, ADEP2 to ADEP 5, Armeniaspirols, phenyl ester compounds (AV126, AV145, AV167, AV170), synthetic peptidomimetic boronate inhibitors (D3, E2, G2, U1), and Hydantoin derivatives were drawn using image identification on the KingDraw software. All ligand structures were saved as .mol files. Sclerotiamide (CID: 10647785), Sclerotiamide P (CID: 170989817), Sclerotiamide L (CID: 170989811), Sclerotiamide N (CID: 170989813), Cyclomarin A (CID: 10772429), Lassomycin (CID: 132528125), Ecumicin (<u>CID: 101894301</u>), Rufomycin B (CID: 76871757), Sclerotiamide B (CID: 146682810), Quercetin (CID: 5280343), Epigallocatechin gallate (EGCG) (CID: 65064), Kaempferol (CID: 5280863), Fisetin (CID: 5281614), Apigenin (CID: 5280443), Luteolin (CID: 5280445) and Genistein (CID: 5280961) were downloaded in the .sdf format from PubChem (https://pubchem.ncbi.nlm.nih.gov/, Accessed 29 July 2024).

### 1.4.3. Ligand ADMET and Toxicity Analysis

The ProTox 3.0 server was used to predict the toxicity of the small ligands to ensure that they were suitable for use in humans (Banerjee *et al*., 2024). The SWISSADME server (http://www.swissadme.ch/; Accessed 30 July 2024) was used to determine the pharmacodynamics of the small ligands through ADME (Adsorption, Distribution, Metabolism, Excretion) analysis (Daina *et al*., 2017). The pkCSM server was used to predict ADMET properties of the large ligands (Pires *et al*., 2015).

### 1.4.4. Molecular Docking studies

The ligands selected for analysis were prepared using the LipPrep module on the Maestro Schrodinger software (Yang *et al*., 2021). Molecular docking was performed using the Autodock vina program integrated on the PyRx software (Ayodele *et al*., 2023). Briefly, the prepared protein was loaded into the PyRx software and made into a macromolecule for docking analysis. The ligands were then loaded into the PyRx software and converted into ligands for docking and saved as .pdbqt files. The grid (Center: X = - 1.1473, Y = -1.1450, Z = -3.1984, Dimensions: X = 93.9220, Y = 105.3030, Z = 107.8333) was set to cover the entirety of ClpK and docking was performed. ClpK-ligand interactions were visualised using Discovery Studio Visualiser 2023 v24.1.0.23298 and PyMol (Schrödinger, 2015).

### 1.4.5. Molecular dynamics simulations using GROMACS

Molecular dynamics (MD) simulations are computational methods used to predict how protein atoms or other organic systems move over time (Hollingsworth and Dror, 2018). MD simulations were performed using GROMACS, an open-source and widely used software package for biomolecular simulation (Abraham *et al*., 2015). The Graphics processing unit (GPU)-accelerated version of GROMACS v2024.2 was installed on a Windows Subsystem for Linux (WSL) 2. Simulations were performed for 100 ns on ClpK alone and selected ClpK-ligand complexes in the presence of either 0.15 M NaCl and 0.05 M MgCl_2_, to assess structural stability and interaction dynamics. The simulation workflow was adapted based on established protocols available on the official GROMACS tutorial website (http://www.mdtutorials.com/gmx/) and is briefly summarised below (Lemkul, 2018).

#### 1.4.5.1. ClpK MD simulation of in the absence of ligands

The refined ClpK model was subjected to MD simulation using GROMACS. The initial structure file (protein.pdb) was processed using the pdb2gmx module to generate the topology, position restraint and post-processed (protein.gro) structure files. The all-atom Optimized Potentials for Liquid Simulations (OPLS) forcefield was used, and solvent model specified as SPC/E. A triclinic simulation box was generated with a minimum distance of 1.0 nm between ClpK and box edges. Solvation was performed using transferable intermolecular potential with 3 points (TIP3P) water model with the predefined SPC216 configuration. The system was then neutralised, and ions were added (either 0.15 M NaCl or 0.05 M MgCl_2_). Energy minimisation was performed using the steepest descent algorithm for 50000 steps to relieve steric clashes and optimize the initial geometry. The system was the equilibrated in two steps: NVT (number of particles, volume, temperature) and NPT (number of particles, pressure, temperature). The 100 ps NVT ensemble was used to stabilise the temperature at 300 K. The 100 ps NPT ensemble was used to stabilise the pressure at 1 bar. A production MD was subsequently performed for 100 ns using a 2 fs time step. The Leap-frog integrator was used to solve the equations of motion and Verlet cut-off scheme was used to evaluate short-ranged non-bonded interactions. Temperature coupling was controlled using the modified Berendsen thermostat, and pressure coupling was controlled using the Parrinello-Rahman barostat. Long-range electrostatic interactions were calculated using the Particle Mesh Ewald (PME) method.

#### 1.4.5.2. ClpK-ligand bound MD simulations

To prepare the ClpK-ligand complexes for MD simulations, ClpK and ligand files were saved as separate files. ClpK structure was processed using the Charmm36-jul2022 force field and solvated using the TIP3P water model. The resulting ClpK_processed.gro file was visualised in PyMol to ensure successful model processing. Ligand preparation was performed using Open Babel and Networkx v2.3. A perl script for ligand topology generation was obtained from the GROMACS tutorial site (http://www.mdtutorials.com/gmx/complex/02_topology.html). The ligand.pdb file was converted to ligand.mol2 using Open Babel. The ligand.mol2 file manually edited using a text editor to ensure consistent naming of the ligand (renamed to “LIG”) across the header and atom records, and the bond order was verified. The edited ligand_fix.mol2 file was submitted to the CHARMM General Force Field server (CGennFF) (https://app.cgenff.com/homepage) to generate topology and parameter files (ligand.str). The generated ligand_ini.pdb file was converted to ligand.gro, which was visualised in PyMol to ensure confirm correct formatting and structure. The processed ClpK and ligand files were then combined using a text editor tool and visualised in PyMol to verify proper merging of the complex. The system topology file (topol.top) was manually updated to include ligand topology and parameters. A triclinic simulation box was defined with a 1.0 nm buffer around the complex, and system was solvated using the TIP3P water model. The solvated system was visualised using PyMol to confirm that the ClpK-ligand complex was centred within the box. The system was then neutralised and either 0.15 M NaCl or 0.05 M MgCl_2_ was added. Visual inspection using PyMol confirmed the presence of ions. Energy minimisation was performed to relieve any steric clashes. Prior to equilibration, the ligand was restrained and temperature coupling groups were defined. A position restrain topology file for the ligand was appended to the end of the system topology file. The index.ndx file was examined to confirm successful definition of ClpK, ligand, and solvent groups. Equilibration parameter files (nvt.mdp, md.mdp and npt.mdp) were adapted from the GROMACS tutorial and modified to reflect the custom groups. NVT equilibration was performed for 100 ps at 300K, followed by NPT equilibration for 100 ps at 1 bar. A 100 ns production MD simulation was then performed to evaluate structural dynamics and protein-ligand complex stability.

### 1.4.6. Molecular dynamic simulation post-run analysis

Following the MD simulations, trajectory post-processing was performed to correct for periodic boundary conditions (PBC), which cause the protein to appear broken or diffused across the unit cell. For the *apo*-ClpK simulation the following analyses were conducted: root mean square deviation (RMSD), root mean square fluctuation (RMSF), potential energy, and radius of gyration (Rg). For the ClpK-ligand complexes, additional analyses were conducted to investigate the impact of ligand binding, these included: RMSD, RMSF, Rg, potential energy, hydrogen bond analysis, Solvent Accessible Surface Area (SASA), and Molecular Mechanics/Generalized Born Surface Area (MM/GBSA) (Bondi, 1964).

## 1.5. Author Contributions

Conceptualization, T.K.; methodology, T.B., and T.K.; data curation T.B.; modelling, T.B., and T.K.; validation, T.B., and T.K.; molecular dynamic simulations, T.B.; formal analysis, T.B. and T.K.; investigation, T.B. and T.K.; protein expression and optimization, T.B.; biophysical characterization, T.B., and T.K. writing—original draft preparation, T.B.; writing—review and editing, T.B., and T.K.; visualization, T.B., and T.K.; supervision, T.K.; project administration, T.K.; All authors have read and agreed to the published version of the manuscript.

## 1.6. Funding

Tehrim Ballim would like to thank Department of Science and Technology-National Research Foundation (DST-NRF), South Africa for the Doctoral Scholarship with Grant Number MND210602605517. Thandeka Khoza also thanks the National Research Foundation Grant number 121275, South African Medical Research Council-SIR grant and University of KwaZulu-Natal for research grants.

## 1.7. Conflicts of Interest

The authors declare no conflict of interest. The funders had no role in the design of the study; in the collection, analyses, or interpretation of data; in the writing of the manuscript, or in the decision to publish the result.

## Notes

### Competing Interest Statement

The authors have declared no competing interest.

## References

Abbas, R., Chakkour, M., Zein El Dine, H., Obaseki, E. F., Obeid, S. T., Jezzini, A., Ghssein, G. & Ezzeddine, Z. 2024. General Overview of Klebsiella pneumonia: Epidemiology and the Role of Siderophores in Its Pathogenicity. Biology, 13, 78.

Abraham, M. J., Murtola, T., Schulz, R., Páll, S., Smith, J. C., Hess, B. & Lindahl, E. 2015. GROMACS: High performance molecular simulations through multi-level parallelism from laptops to supercomputers. SoftwareX, 1-2, 19-25.

Afriza, D., Suriyah, W. H. & Ichwan, S. J. A. 2018. In silico analysis of molecular interactions between the anti-apoptotic protein survivin and dentatin, nordentatin, and quercetin. Journal of Physics: Conference Series, 1073, 032001.

Ajao, A., Kannan, M., Yakubu, S., Vj, U. & Jb, A. 2012. Homology modeling, simulation and molecular docking studies of catechol-2, 3-Dioxygenase from Burkholderia cepacia: Involved in degradation of Petroleum hydrocarbons. Bioinformation, 8, 848–54.

Alazmi, M. & Motwalli, O. 2021. In silico virtual screening, characterization, docking and molecular dynamics studies of crucial SARS-CoV-2 proteins. J Biomol Struct Dyn, 39, 6761–6771.

Ali, A., Mir, G. J., Ayaz, A., Maqbool, I., Ahmad, S. B., Mushtaq, S., Khan, A., Mir, T. M. & Rehman, M. U. 2023. In silico analysis and molecular docking studies of natural compounds of Withania somnifera against bovine NLRP9. Journal of Molecular Modeling, 29, 171.

Ayodele, P., Bamigbade, A., Bamigbade, O., Adeniyi, I., Tachin, E., Seweje, A. & Farohunbi, S. 2023. Illustrated Procedure to Perform Molecular Docking Using PyRx and Biovia Discovery Studio Visualizer: A Case Study of 10kt With Atropine. Progress in Drug Discovery & Biomedical Science, 6, 1–32.

Bagley, S. T. 1985. Habitat Association of Klebsiella Species. Infection Control, 6, 52–58.

Banerjee, P., Kemmler, E., Dunkel, M. & Preissner, R. 2024. ProTox 3.0: a webserver for the prediction of toxicity of chemicals. Nucleic Acids Research, 52, W513–W520.

Berg, J. M., Tymoczko, J. L. & Stryer, L. 2007. Biochemistry (loose-leaf), Macmillan.

Bhandari, V., Wong, K. S., Zhou, J. L., Mabanglo, M. F., Batey, R. A. & Houry, W. A. 2018. The Role of ClpP Protease in Bacterial Pathogenesis and Human Diseases. ACS Chem Biol, 13, 1413–1425.

Bojer, M. S., Hammerum, A. M., Jørgensen, S. L., Hansen, F., Olsen, S. S., Krogfelt, K. A. & Struve, C. 2012. Concurrent emergence of multidrug resistance and heat resistance by CTX-M-15-encoding conjugative plasmids in lebsiella pneumoniae. APMIS, 120, 699–705.

Bojer, M. S., Struve, C., Ingmer, H., Hansen, D. S. & Krogfelt, K. A. 2010. Heat resistance mediated by a new plasmid encoded Clp ATPase, ClpK, as a possible novel mechanism for nosocomial persistence of Klebsiella pneumoniae. PLoS One, 5, e15467.

Bondi, A. 1964. van der Waals Volumes and Radii. The Journal of Physical Chemistry, 68, 441–451.

Borjian Boroujeni, M., Shahbazi Dastjerdeh, M., Shokrgozar, M., Rahimi, H. & Omidinia, E. 2021. Computational driven molecular dynamics simulation of keratinocyte growth factor behavior at different pH conditions. Informatics in Medicine Unlocked, 23, 100514.

Bouchnak, I. & Van Wijk, K. J. 2021. Structure, function, and substrates of Clp AAA+ protease systems in cyanobacteria, plastids, and apicoplasts: A&#xa0;comparative analysis. Journal of Biological Chemistry, 296.

Brylinski, M. 2018. Aromatic interactions at the ligand-protein interface: Implications for the development of docking scoring functions. Chem Biol Drug Des, 91, 380–390.

Carroni, M., Franke, K. B., Maurer, M., Jäger, J., Hantke, I., Gloge, F., Linder, D., Gremer, S., Turgay, K., Bukau, B. & Mogk, A. 2017. Regulatory coiled-coil domains promote head-to-head assemblies of AAA+ chaperones essential for tunable activity control. eLife, 6, e30120.

Chaturvedi, U. C. & Shrivastava, R. 2005. Interaction of viral proteins with metal ions: role in maintaining the structure and functions of viruses. FEMS Immunol Med Microbiol, 43, 105–14.

Choules, M. P., Wolf, N. M., Lee, H., Anderson, J. R., Grzelak, E. M., Wang, Y., Ma, R., Gao, W., Mcalpine, J. B., Jin, Y. Y., Cheng, J., Lee, H., Suh, J. W., Duc, N. M., Paik, S., Choe, J. H., Jo, E. K., Chang, C. L., Lee, J. S., Jaki, B. U., Pauli, G. F., Franzblau, S. G. & Cho, S. 2019. Rufomycin Targets ClpC1 Proteolysis in Mycobacterium tuberculosis and M. abscessus. Antimicrob Agents Chemother, 63.

Colovos, C. & Yeates, T. O. 1993. Verification of protein structures: patterns of nonbonded atomic interactions. Protein Sci, 2, 1511–9.

Culp, E. & Wright, G. D. 2017. Bacterial proteases, untapped antimicrobial drug targets. The Journal of Antibiotics, 70, 366–377.

Daina, A., Michielin, O. & Zoete, V. 2017. SwissADME: a free web tool to evaluate pharmacokinetics, drug-likeness and medicinal chemistry friendliness of small molecules. Sci Rep, 7, 42717.

Daya, T., Jeje, O., Maake, R., Chinyere, A., Khoza, T. & Achilonu, I. 2022. Expression, Purification, and Biophysical Characterization of Klebsiella Pneumoniae Nicotinate Nucleotide Adenylyltransferase. The Protein Journal, 41, 1–16.

De Campos, L. J., Seleem, M. A., Feng, J., Pires De Oliveira, K. M., De Andrade Dos Santos, J. V., Hayer, S., Clayton, J. B., Kathi, S., Fisher, D. J., Ouellette, S. P. & Conda-Sheridan, M. 2023. Design, Biological Evaluation, and Computer-Aided Analysis of Dihydrothiazepines as Selective Antichlamydial Agents. J Med Chem, 66, 2116–2142.

Ferenczy, G. G. & Kellermayer, M. 2022. Contribution of hydrophobic interactions to protein mechanical stability. Comput Struct Biotechnol J, 20, 1946–1956.

Ferrari, S., Costi, P. M. & Wade, R. C. 2003. Inhibitor Specificity via Protein Dynamics: Insights from the Design of Antibacterial Agents Targeted Against Thymidylate Synthase. Chemistry & Biology, 10, 1183–1193.

Fetzer, C., Korotkov, V. S. & Sieber, S. A. 2019. Hydantoin analogs inhibit the fully assembled ClpXP protease without affecting the individual peptidase and chaperone domains. Organic & Biomolecular Chemistry, 17, 7124–7127.

Fetzer, C., Korotkov, V. S., Thänert, R., Lee, K. M., Neuenschwander, M., Von Kries, J. P., Medina, E. & Sieber, S. A. 2017. A Chemical Disruptor of the ClpX Chaperone Complex Attenuates the Virulence of Multidrug-Resistant Staphylococcus aureus. Angewandte Chemie International Edition, 56, 15746–15750.

Gavrish, E., Sit, C. S., Cao, S., Kandror, O., Spoering, A., Peoples, A., Ling, L., Fetterman, A., Hughes, D., Bissell, A., Torrey, H., Akopian, T., Mueller, A., Epstein, S., Goldberg, A., Clardy, J. & Lewis, K. 2014. Lassomycin, a ribosomally synthesized cyclic peptide, kills mycobacterium tuberculosis by targeting the ATP-dependent protease ClpC1P1P2. Chem Biol, 21, 509–518.

Gersch, M., Famulla, K., Dahmen, M., Göbl, C., Malik, I., Richter, K., Korotkov, V. S., Sass, P., Rübsamen-Schaeff, H., Madl, T., Brötz-Oesterhelt, H. & Sieber, S. A. 2015. AAA+ chaperones and acyldepsipeptides activate the ClpP protease via conformational control. Nature Communications, 6, 6320.

Hackl, M. W., Lakemeyer, M., Dahmen, M., Glaser, M., Pahl, A., Lorenz-Baath, K., Menzel, T., Sievers, S., Böttcher, T., Antes, I., Waldmann, H. & Sieber, S. A. 2015. Phenyl Esters Are Potent Inhibitors of Caseinolytic Protease P and Reveal a Stereogenic Switch for Deoligomerization. J Am Chem Soc, 137, 8475–83.

Han, S., Mei, L., Quach, T., Porter, C. & Trevaskis, N. 2021. Lipophilic Conjugates of Drugs: A Tool to Improve Drug Pharmacokinetic and Therapeutic Profiles. Pharm Res, 38, 1497–1518.

Hollingsworth, S. A. & Dror, R. O. 2018. Molecular Dynamics Simulation for All. Neuron, 99, 1129–1143.

Ishikawa, F., Homma, M., Tanabe, G. & Uchihashi, T. 2024. Protein degradation by a component of the chaperonin-linked protease ClpP. Genes to Cells, n/a.

Jernigan, R. L., Khade, P., Kumar, A. & Kloczkowski, A. 2022. Using Surface Hydrophobicity Together with Empirical Potentials to Identify Protein-Protein Binding Sites: Application to the Interactions of E-cadherins. Methods Mol Biol, 2340, 41–50.

Jumper, J., Evans, R., Pritzel, A., Green, T., Figurnov, M., Ronneberger, O., Tunyasuvunakool, K., Bates, R., Žídek, A., Potapenko, A., Bridgland, A., Meyer, C., Kohl, S. a. A., Ballard, A. J., Cowie, A., Romera-Paredes, B., Nikolov, S., Jain, R., Adler, J., Back, T., Petersen, S., Reiman, D., Clancy, E., Zielinski, M., Steinegger, M., Pacholska, M., Berghammer, T., Bodenstein, S., Silver, D., Vinyals, O., Senior, A. W., Kavukcuoglu, K., Kohli, P. & Hassabis, D. 2021. Highly accurate protein structure prediction with AlphaFold. Nature, 596, 583–589.

Khan, N., Syed, D. N., Ahmad, N. & Mukhtar, H. 2013. Fisetin: a dietary antioxidant for health promotion. Antioxid Redox Signal, 19, 151–62.

Kirstein, J., Hoffmann, A., Lilie, H., Schmidt, R., Rübsamen-Waigmann, H., Brötz-Oesterhelt, H., Mogk, A. & Turgay, K. 2009. The antibiotic ADEP reprogrammes ClpP, switching it from a regulated to an uncontrolled protease. EMBO Mol Med, 1, 37–49.

Ko, J., Park, H., Heo, L. & Seok, C. 2012. GalaxyWEB server for protein structure prediction and refinement. Nucleic Acids Research, 40, W294–W297.

Labana, P., Dornan, M. H., Lafrenière, M., Czarny, T. L., Brown, E. D., Pezacki, J. P. & Boddy, C. N. 2021. Armeniaspirols inhibit the AAA+ proteases ClpXP and ClpYQ leading to cell division arrest in Gram-positive bacteria. Cell Chemical Biology, 28, 1703–1715.e11.

Lairson, L. L., Henrissat, B., Davies, G. J. & Withers, S. G. 2008. Glycosyltransferases: Structures, Functions, and Mechanisms. Annual Review of Biochemistry, 77, 521–555.

Lavey, N. P., Coker, J. A., Ruben, E. A. & Duerfeldt, A. S. 2016. Sclerotiamide: The First Non-Peptide-Based Natural Product Activator of Bacterial Caseinolytic Protease P. J Nat Prod, 79, 1193–7.

Lemkul, J. A. 2018. From Proteins to Perturbed Hamiltonians: A Suite of Tutorials for the GROMACS-2018 Molecular Simulation Package [Article v1.0]. Living Journal of Computational Molecular Science, 1, 5068.

Li, Y., Kumar, S., Zhang, L., Wu, H. & Wu, H. 2023. Characteristics of antibiotic resistance mechanisms and genes of Klebsiella pneumoniae. Open Medicine, 18.

Madushanka, A., Moura, R. T., Verma, N. & Kraka, E. 2023. Quantum Mechanical Assessment of Protein–Ligand Hydrogen Bond Strength Patterns: Insights from Semiempirical Tight-Binding and Local Vibrational Mode Theory. International Journal of Molecular Sciences, 24, 6311.

Martin, Y. 2005. A Bioavailability Score. Journal of medicinal chemistry, 48, 3164–70.

Mfeka, M., Onisuru, O., Pandian, R. P., Sayed, Y., Khoza, T. & Achilonu, I. 2025. Crystal enigma: Understanding diverse protein conformational dynamics, ligand selectivity and interaction in multi-space group crystals using computational modelling. Results in Chemistry, 16, 102288.

Molaakbari, E., Aallae, M. R., Golestanifar, F., Garakani-Nejad, Z., Khosravi, A., Rezapour, M., Eshaghi Malekshah, R., Ghomi, M. & Ren, G. 2024. Insilico assessment of hesperidin on SARS-CoV-2 main protease and RNA polymerase: Molecular docking and dynamics simulation approach. Biochemistry and Biophysics Reports, 39, 101804.

Moreno-Cinos, C., Goossens, K., Salado, I. G., Van Der Veken, P., De Winter, H. & Augustyns, K. 2019. ClpP Protease, a Promising Antimicrobial Target. Int J Mol Sci, 20.

Motiwala, T., Akumadu, B. O., Zuma, S., Mfeka, M. S., Chen, W., Achilonu, I., Syed, K. & Khoza, T. 2021. Caseinolytic Proteins (Clp) in the Genus Klebsiella: Special Focus on ClpK. Molecules, 27.

Motiwala, T., Mthethwa, Q., Achilonu, I. & Khoza, T. 2022. ESKAPE Pathogens: Looking at Clp ATPases as Potential Drug Targets. Antibiotics, 11, 1218.

Motiwala, T., Nyide, B. & Khoza, T. 2025. Molecular dynamic simulations to assess the structural variability of ClpV from Enterobacter cloacae. *Frontiers in Bioinformatics*, Volume 5–2025.

Mulani, M. S., Kamble, E. E., Kumkar, S. N., Tawre, M. S. & Pardesi, K. R. 2019. Emerging Strategies to Combat ESKAPE Pathogens in the Era of Antimicrobial Resistance: A Review. Frontiers in Microbiology, 10.

Pires, D. E. V., Blundell, T. L. & Ascher, D. B. 2015. pkCSM: Predicting Small-Molecule Pharmacokinetic and Toxicity Properties Using Graph-Based Signatures. Journal of Medicinal Chemistry, 58, 4066–4072.

Plotniece, A., Sobolev, A., Supuran, C. T., Carta, F., Björkling, F., Franzyk, H., Yli-Kauhaluoma, J., Augustyns, K., Cos, P., De Vooght, L., Govaerts, M., Aizawa, J., Tammela, P. & Žalubovskis, R. 2023. Selected strategies to fight pathogenic bacteria. J Enzyme Inhib Med Chem, 38, 2155816.

Podschun, R. & Ullmann, U. 1998. Klebsiella spp. as nosocomial pathogens: epidemiology, taxonomy, typing methods, and pathogenicity factors. Clin Microbiol Rev, 11, 589–603.

Prestinaci, F., Pezzotti, P. & Pantosti, A. 2015. Antimicrobial resistance: a global multifaceted phenomenon. Pathog Glob Health, 109, 309–18.

Rei Liao, J.-Y. & Van Wijk, K. J. 2019. Discovery of AAA+ Protease Substrates through Trapping Approaches. Trends in Biochemical Sciences, 44, 528–545.

Schauss, J., Kundu, A., Fingerhut, B. P. & Elsaesser, T. 2021. Magnesium Contact Ions Stabilize the Tertiary Structure of Transfer RNA: Electrostatics Mapped by Two-Dimensional Infrared Spectra and Theoretical Simulations. J Phys Chem B, 125, 740–747.

Schmitt, E. K., Riwanto, M., Sambandamurthy, V., Roggo, S., Miault, C., Zwingelstein, C., Krastel, P., Noble, C., Beer, D., Rao, S. P. S., Au, M., Niyomrattanakit, P., Lim, V., Zheng, J., Jeffery, D., Pethe, K. & Camacho, L. R. 2011. The Natural Product Cyclomarin Kills Mycobacterium Tuberculosis by Targeting the ClpC1 Subunit of the Caseinolytic Protease. Angewandte Chemie International Edition, 50, 5889–5891.

Schönichen, A., Webb, B. A., Jacobson, M. P. & Barber, D. L. 2013. Considering protonation as a posttranslational modification regulating protein structure and function. Annu Rev Biophys, 42, 289–314.

Schrödinger, L. L. C. 2015. The PyMOL Molecular Graphics System, Version 2.5.

Schwede, T., Kopp, J., Guex, N. & Peitsch, M. C. 2003. SWISS-MODEL: An automated protein homology-modeling server. Nucleic Acids Res, 31, 3381–5.

Seraphim, T. V. & Houry, W. A. 2020. AAA+ proteins. Current Biology, 30, R251–R257.

Shamsara, J. 2018. Correlation between Virtual Screening Performance and Binding Site Descriptors of Protein Targets. Int J Med Chem, 2018, 3829307.

Shao, D., Zhang, Q., Xu, P. & Jiang, Z. 2022. Effects of the Temperature and Salt Concentration on the Structural Characteristics of the Protein (PDB Code 1BBL). Polymers, 14, 2134.

Shrivastava, S. R., Shrivastava, P. S. & Ramasamy, J. 2018. World health organization releases global priority list of antibiotic-resistant bacteria to guide research, discovery, and development of new antibiotics. Journal of Medical Society, 32, 76–77.

Silakari, O. & Singh, P. K. 2021. Chapter 14 - ADMET tools: Prediction and assessment of chemical ADMET properties of NCEs. In: Silakari, O. & Singh, P. K. (eds.) Concepts and Experimental Protocols of Modelling and Informatics in Drug Design. Academic Press.

Stratiichuk, R., Melnychenko, M., Koleiev, I., Voitsitskyi, T., Husak, V., Shevchuk, N., Ostrovsky, Z., Bdzhola, V., Yesylevskyy, S., Starosyla, S. & Nafiiev, A. 2025. Leveraging large language models for literature-driven prioritization of protein binding pockets. Bioinformatics, 41.

Varma, M. V. S., Perumal, O. P. & Panchagnula, R. 2006. Functional role of P-glycoprotein in limiting peroral drug absorption: optimizing drug delivery. Current Opinion in Chemical Biology, 10, 367–373.

Vass, R. H. & Chien, P. 2016. Two ways to skin a cat: acyldepsipeptides antibiotics can kill bacteria through activation or inhibition of ClpP activity. Mol Microbiol, 101, 183–5.

Volkamer, A., Griewel, A., Grombacher, T. & Rarey, M. 2010. Analyzing the Topology of Active Sites: On the Prediction of Pockets and Subpockets. Journal of Chemical Information and Modeling, 50, 2041–2052.

Waterhouse, A., Bertoni, M., Bienert, S., Studer, G., Tauriello, G., Gumienny, R., Heer, F. T., De beer, T. A p., Rempfer, C., Bordoli, L., Lepore, R. & Schwede, T. 2018. SWISS-MODEL: homology modelling of protein structures and complexes. Nucleic Acids Research, 46, W296–W303.

Wiederstein, M. & Sippl, M. J. 2007. ProSA-web: interactive web service for the recognition of errors in three-dimensional structures of proteins. Nucleic Acids Research, 35, W407–W410.

Willard, L., Ranjan, A., Zhang, H., Monzavi, H., Boyko, R. F., Sykes, B. D. & Wishart, D. S. 2003. VADAR: a web server for quantitative evaluation of protein structure quality. Nucleic Acids Res, 31, 3316–9.

Williams, C. J., Headd, J. J., Moriarty, N. W., Prisant, M. G., Videau, L. L., Deis, L. N., Verma, V., Keedy, D. A., Hintze, B. J., Chen, V. B., Jain, S., Lewis, S. M., Arendall, W. B., 3rd, Snoeyink, J., Adams, P. D., Lovell, S. C., Richardson, J. S. & Richardson, D. C. 2018. MolProbity: More and better reference data for improved all-atom structure validation. Protein Sci, 27, 293–315.

Xuan, J., Feng, W., Wang, J., Wang, R., Zhang, B., Bo, L., Chen, Z.-S., Yang, H. & Sun, L. 2023. Antimicrobial peptides for combating drug-resistant bacterial infections. Drug Resistance Updates, 68, 100954.

Yang, Y., Yao, K., Repasky, M. P., Leswing, K., Abel, R., Shoichet, B. K. & Jerome, S. V. 2021. Efficient Exploration of Chemical Space with Docking and Deep Learning. Journal of Chemical Theory and Computation, 17, 7106–7119.

Zanini, C., Giribaldi, G., Mandili, G., Carta, F., Crescenzio, N., Bisaro, B., Doria, A., Foglia, L., Di Montezemolo, L. C., Timeus, F. & Turrini, F. 2007. Inhibition of heat shock proteins (HSP) expression by quercetin and differential doxorubicin sensitization in neuroblastoma and Ewing’s sarcoma cell lines. J Neurochem, 103, 1344–54.

Zhao, H. & Huang, D. 2011. Hydrogen Bonding Penalty upon Ligand Binding. PLOS ONE, 6, e19923.

Zininga, T., Ramatsui, L., Makhado, P. B., Makumire, S., Achilinou, I., Hoppe, H., Dirr, H. & Shonhai, A. 2017. (-)-Epigallocatechin-3-Gallate Inhibits the Chaperone Activity of Plasmodium falciparum Hsp70 Chaperones and Abrogates Their Association with Functional Partners. Molecules, 22.

